# The *W*-index: a novel tool to evaluate gender equity in STEM research

**DOI:** 10.1101/2025.10.29.685415

**Authors:** E. Penelope Holland, Jalene M. LaMontagne, V. Bala Chaudhary, Suw Charman-Anderson, Lindsey Gillson, Thorunn Helgason, Angela Lipscomb, Elva J. H. Robinson, Alex James

## Abstract

Gender inequities in scientific research persist, from citation indices to collaboration networks and funding success. Current estimates for achieving equal representation remain in timescales of decades to centuries. To help individuals and groups improve their gender representation in a shorter timeframe, we present the *W*-index, a simple metric to evaluate gender ratios, and specifically the representation of women, in research collaborations and other aspects of scientific life. Here it is applied to three case studies: internal collaborations derived from six years of research outputs across a university, co-author groups from 60 years of publications across a journal collection, and supervisor-student relationships over 60 years. Despite contrasting sources, these datasets show common features: *W*-index increases as the proportion of women increases *W*-index also increases with age or career stage, and women have a consistently higher *W*-index compared to men. We finish with reflections and recommendations for individuals, organisations, and funding bodies to action positive and timely change towards gender equity.

## Main

The proportion of women in science, technology, engineering, and mathematics (STEM) disciplines is still far below parity with men^1,2^ despite strong evidence that diversity improves outcomes^3^. Even in disciplines where undergraduate intake has been majority women for 30 years or more, this still does not translate into a corresponding majority in senior positions or publications^4^ and at the current rate of progress, the gender gap is likely to persist for generations^5,6^. Limited recruitment, retention and success of women and other minorities in science is often framed as a ‘leaky pipeline’, implying women are passively leaking out of the system rather than the “hostile obstacle course”^7^ of systemic gender bias in STEM communities.

Efforts to understand the problem include extensive research on gendered differences in student-supervisor relationships^8^, research presentations^9^, service roles^10,11^ and research evaluation^12,13^, but we now need action rather than words. The cultural transformation required to shift individuals, organisations and funding bodies towards an equitable and just community is a huge task. The evidence shows that suggestions for positive change are overwhelmingly taken up by those already disadvantaged by the system, rather than by everyone^14^. We need to share the mental and emotional load, as well as the tangible work and outputs of science.

To create this change, we introduce the *W*-index (pronounced ‘windex’), a metric so simple it can be used on the spot by anyone in almost any situation. The *W*-index allows individuals working in science to reflect on and articulate the gender composition of their community and collaborators in an instant, to empower individuals to be intentional as they progress through their careers. We show how the W-index varies with age, discipline and gender in three very different situations. We then provide recommendations for how it can be used across all aspects of an academic or scientific career.

### The *W*-index

Very simply, an individual’s *W*-index is the proportion of their collaborators, co-workers or students who are women. This could be co-authors on a single paper or across multiple papers, the research students they have supervised, undergraduates they teach, or colleagues with whom they apply for funding. A person may have different *W*-indices in different situations. Whether applied to co-author networks, supervisor-student relationships or conference panel discussion members, the core question remains the same: are women equitably represented in an individual’s projects, publications or workspaces? The calculation excludes the focal person in question, so the null expectation is that, on average, men and women can have equal *W*-indices.

In a world with stable, equal representation of men and women, all *W*-indices, in any context, should tend towards one half: collaborators, co-workers and students are equally likely to be men as women. Since women are not currently represented equally in all STEM fields, we expect most scientists to have a *W-*index of less than 0.5, but these indices should show an increase over recent decades reflecting the increasing proportion of women in the STEM professional population. If we are to transition towards equity of gender representation and inclusion, we need a clear and deliberate direction toward a lifetime average *W*-index of 0.5.

We present three examples of the *W*-index in action: (1) research collaborations across all disciplines within a single institution (2012-2018); (2) co-authorship in a research and management science journal collection over 55 years (1952-2016); (3) research degree student-supervisor relationships in mathematics over 60 years (1960-2019). We analysed *W*- index using generalised linear mixed-effects models^15-17^ of individual collaborations as a function of (a) gender, (b) career stage or author age, and (c) representation (the proportion of women in a field), and the best model was selected using AIC^18^.

### Research collaborations in a single institution

The internal research output database of the University of Canterbury, Aotearoa New Zealand (UC) contains all academic research outputs (publications, books, technical reports, etc.) published over a six-year period (2012-2018) that have at least one UC author. It includes publication date and the age, gender and academic department for each UC author. Non-UC authors are not described in detail. The dataset contains 8,441 unique research outputs for 1,304 unique authors, combined into over 20,000 author-output pairs, i.e. most authors have more than one publication and some publications have more than one UC author. Authors are from 29 academic departments and 41% are women. Single author outputs, i.e. one UC author, were excluded. Gender was assigned using university personnel records. NZ allows transgender individuals to self-report their gender and there were no non-binary individuals in the records. Authors with unknown or unreported gender were excluded.

*W*-index for each individual was calculated as the proportion of UC co-authors on a publication who were women averaged across all the authors publications. Publications were equally weighted regardless of the number of co-authors.

The mean *W*-index for women at UC was 0.47, i.e. almost half of their co-authors on any paper were women, with high variance across individuals (interquartile range, IQR, of 0-1.0), while for men, the mean *W*-index was lower at 0.23 and the variance was smaller (IQR of 0-0.5). Individuals in departments with more women (i.e., Psychology, Arts) have a higher *W*-index (Figure 1), equating to an increased *W*-index of about 0.07 for every 10% shift in gender balance (linear regression on age, gender and departmental gender balance; see Supplementary Information). In a department with equal numbers of men and women, the average *W*-index of all individuals increases by 0.02 per ten years of age, but the average *W*- index of an early career woman (age 30) is 0.16 higher than an equivalent man, and the gap narrows to 0.1 at age 50. The average *W*-index of an individual is almost always lower than the proportion of women in their department.

**Figure 1.**
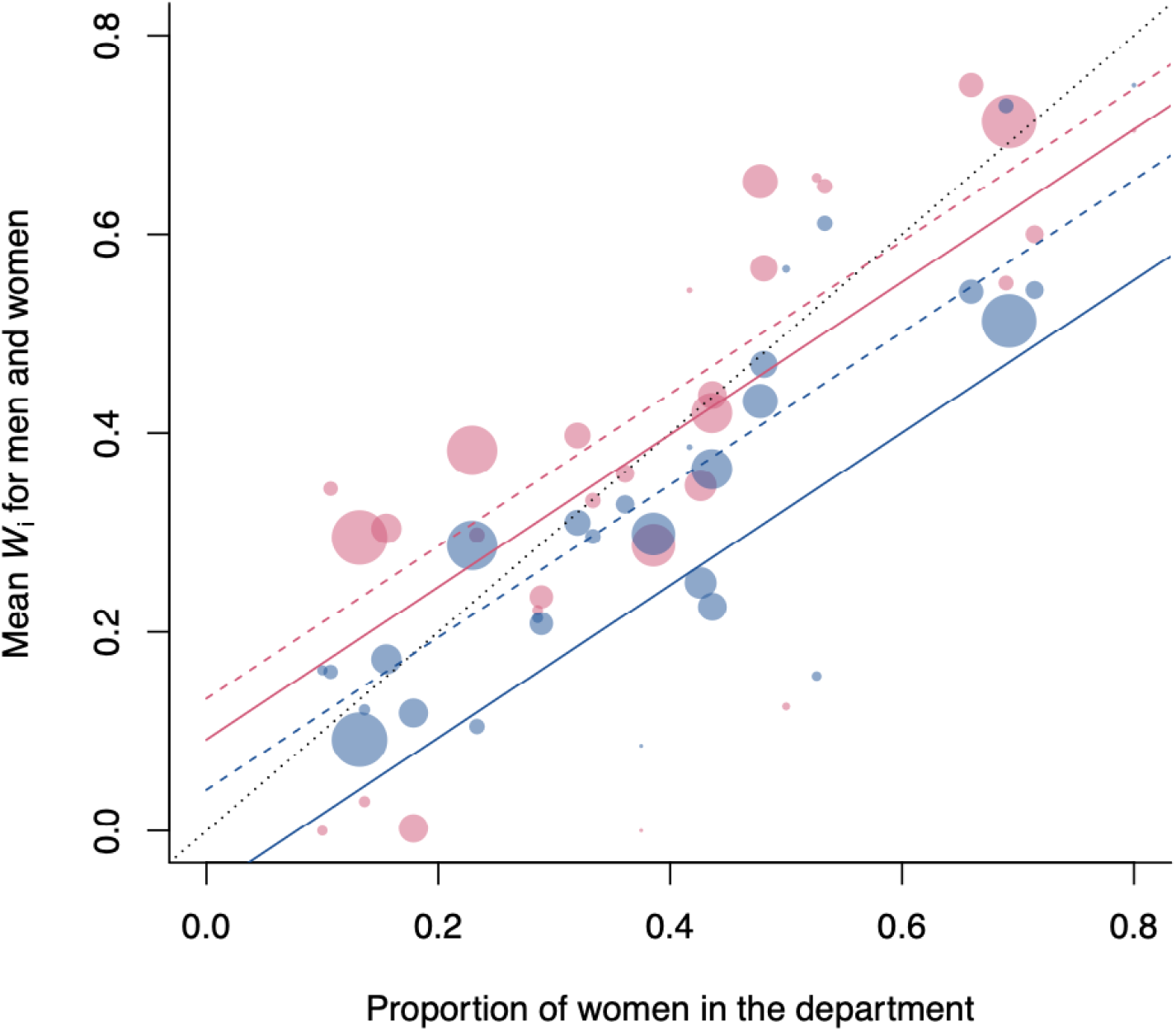
Women, particularly older women, have a higher *W*-index than men. The average *W*-index of men (blue circles) and women (pink circles) in each department increases with increasing department gender balance (a proxy for women in the discipline). Circle size is proportional to department size. *W*-index increases with increasing proportion of women in the department, and with age (predicted *W*-index for authors age 30 in solid lines; dashed lines for age 50), but largely remains below the 1:1 line (dotted line). For example, in a 50:50 gender-balanced department, a 30-year-old man has, on average, only 32% woman co-authors on a given output, and a 50-year-old man has 42%. In contrast, 30- and 50-year-old women in the same department have 48 and 52% women co-authors respectively.

### Co-authorships in a journal collection

The INFORMS dataset^19^, published through the Institute for Operations Research and Management Sciences, contains records of over 17,000 individual authors linked to over 14,000 multi-author publications, resulting in approximately 37,000 author-publication pairs, spanning 55 years (1952 to 2016). Pairs where the author gender was unknown were excluded. Overall, 16% of authors are women, and in 13% of author-publication pairs the author is a woman. Authors were assigned early career status on outputs published within 10 years of the author’s first appearance in the dataset, and established career status otherwise. A threshold of 5 or 15 years makes little difference to the results (see Supplementary Information).

*W*-index for each author-publication pair was calculated as the mean proportion of co-authors on the publication who were women, excluding the author in question. To find an author’s average *W*-index publications from that author were equally weighted regardless of the number of co-authors.

The mean *W*-index for women was 0.22 (IQR = 0.0-0.4) and for men the mean was 0.12 and the variation was also smaller (IQR = 0.0-0.14). The proportion of authorships by women increases from almost zero in 1952 to 0.15 over the timespan of the data, and women authorships are predominantly found in the later years of the data period when there were proportionally more women. The *W*-index for both men and women increases with the representation of women over time (Figure 2, linear regression of *W*-index for late (dashed lines) and early (solid lines) stage authors accounting for proportion of women authors in the dataset in the publication year). Men publish in teams with gender balance roughly in line with the proportion of women in the dataset that year but women’s *W*-indices are consistently higher and increase with seniority, whereas men’s *W-*indices change very little from early to established career.

**Figure 2.**
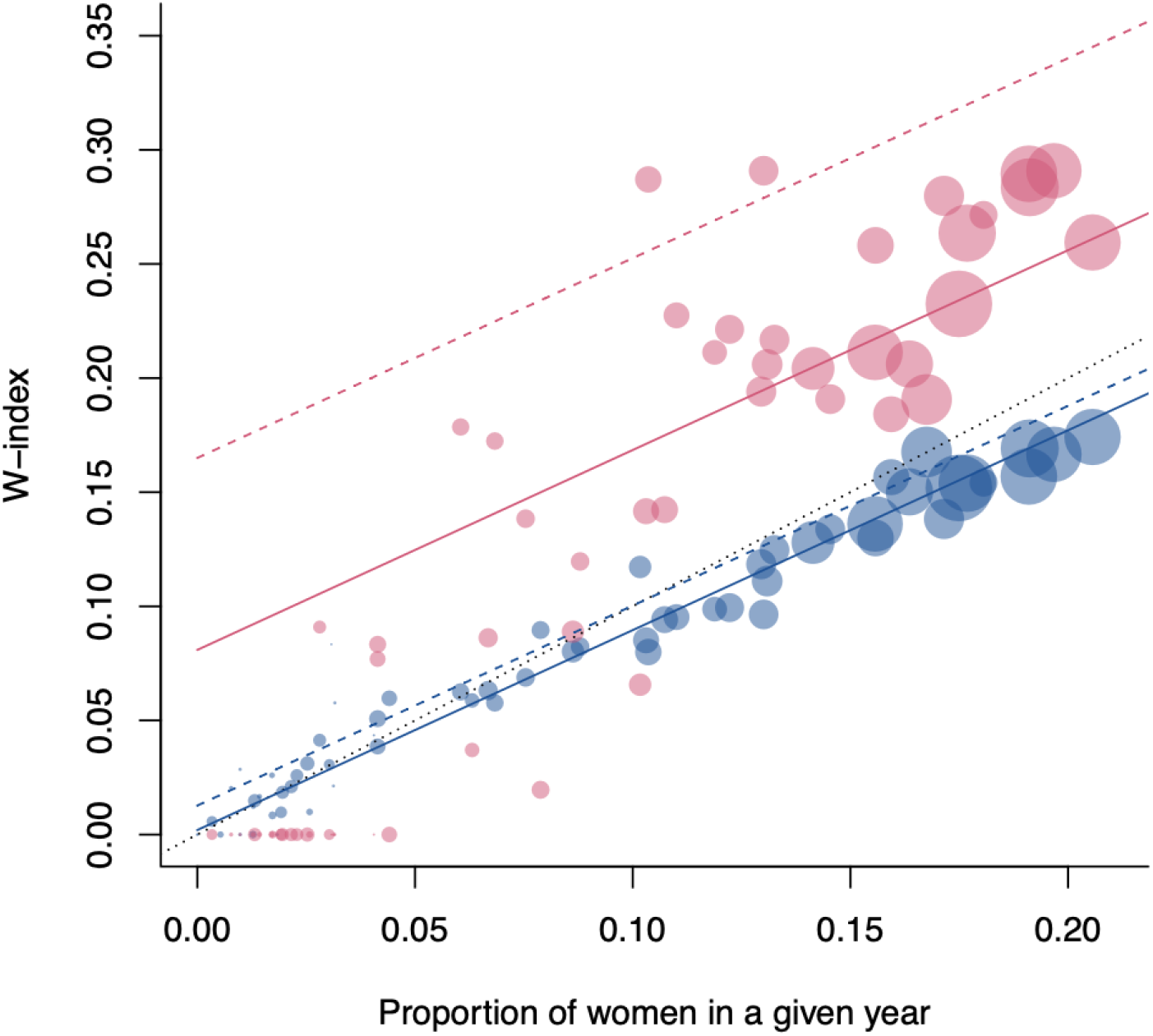
Women are poorly represented overall, and in collaborations. Women, especially late career women, have a higher *W*-index than men. Mean *W*-index for men (blue) and women (pink) in the INFORMS journal dataset by year, against the proportion of authors publishing in a given year that are women. Circle size represents total number of author-publication pairs that year. Lines show the expected (model) *W*-index for early career (solid) or established career (dashed) authors, and the 1:1 line (black dots).

### Student-supervisor relationships for research degrees

The Maths Genealogy Project (MGP) is a citizen science database of mathematical science research degrees. Any individual can enter the details of their research degree in any area of mathematical science. The data includes the names of the student and supervisors and the year and institution of the degree. After names were gendered (see Supplementary Information) this gave 77,985 student-supervisor pairs from 19,354 supervisors. Supervisor career stage for each supervisor-student pair was assigned by years since first supervision by the supervisor in the dataset, with a 10-year threshold for early vs established.

*W*-index for each supervisor-student pair was simply zero or one depending on the gender of the student. A supervisor’s average lifetime *W*-index was the proportion of students they had supervised who were female. The mean lifetime *W*-index for women supervisors was 0.34, i.e. on average one third of students supervised by women were women, (IQR = 0.0-0.67) and for men, the mean was 0.17 (IQR = 0.0-0.25). The proportion of students who are women increased from 4.3% in 1960 to 23.9% in 2000 but did not rise substantially in the twenty years following. The *W*-index for both men and women supervisors increases with the representation of women (Figure 3). While men supervise students roughly in line with the proportion of women in the student population that year, women’s *W*-indices are consistently higher. Established supervisors who are men have a higher *W*-index than their early career counterparts, whereas the opposite is true for women.

**Figure 3.**
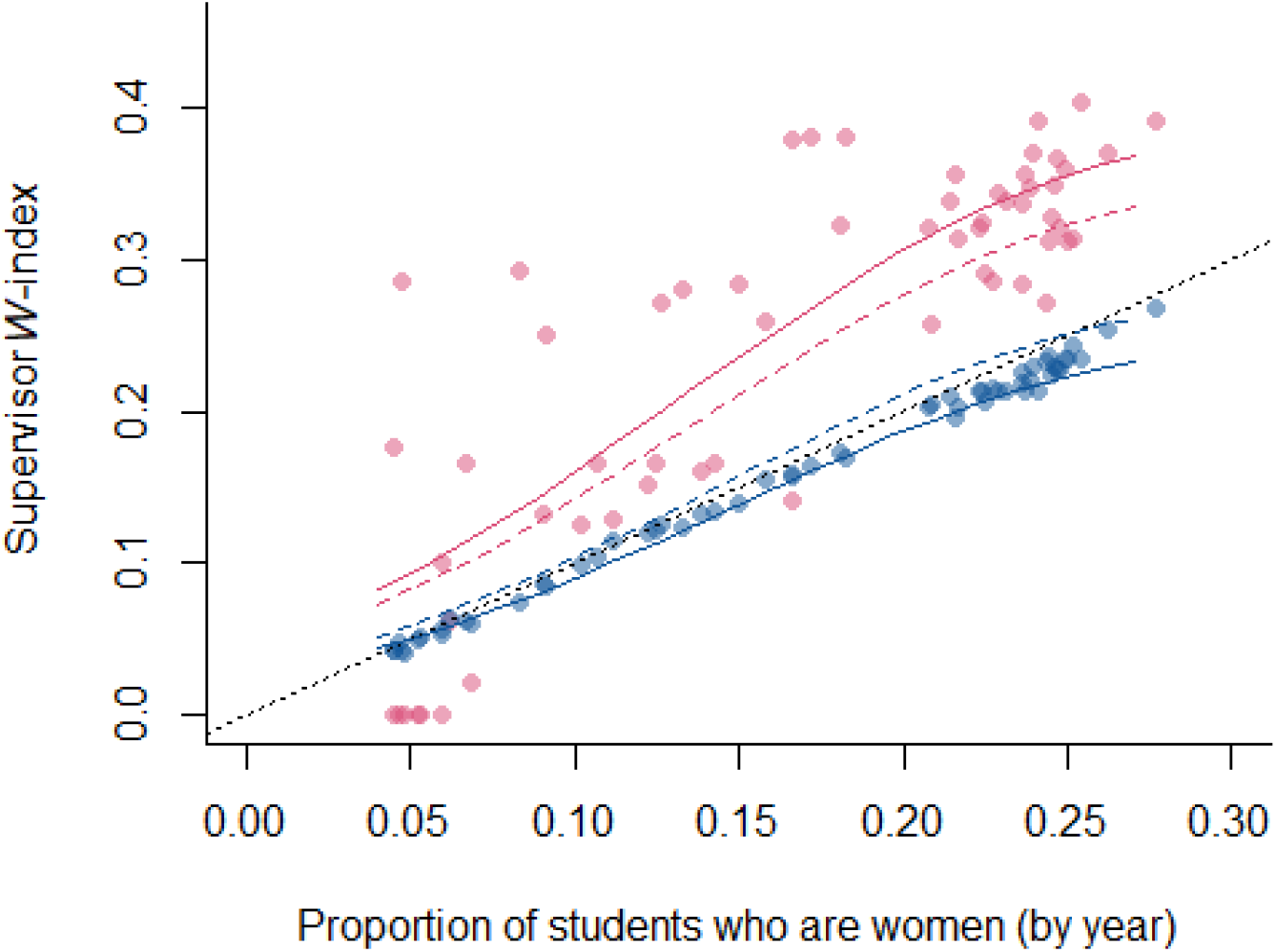
Women are poorly represented as trainees overall, and women research supervisors have a higher *W*-index than men. The average proportion of students who are women supervised by men (blue circles) and women (pink circles) supervisors in each year against the proportion of students who are women each year. Mixed-effects logistic regression indicates that the *W*-index of all supervisors increases as overall representation of women in the student body increases (lines). For men (blue lines) this tracks below the 1:1 line (black dots), for women above. Established career women (dashed pink line) consistently supervise slightly fewer women than early career women (solid pink line), whereas men increase *W*- index slightly in late career (dashed blue line) compared to early career (solid blue line) but still predominantly below the 1:1 line.

### Towards a new norm

Despite the contrasting sources, all three data sets show three common features. *W*-index increases with higher representation of women, which is expected. Women also have a consistently higher *W*-index compared to men, and age/career stage have a significant effect on *W*-index.

The *W*-index of men tracks below the proportion of women available to work with (Figures 1 and 2) or to supervise (Figure 3), whereas women have *W*-indices that are 0.1 or more higher than those of men. Research in chemistry indicates that students working with advisors of the same gender as themselves are likely to be more productive during their postgraduate research student period, while women working with women are also more likely to progress into academic (faculty) roles^8^. Gender homophily, the tendency to work with people of the same gender, is well-known in academic publications^20^ and networks^21^. The *W-*index generalises and quantifies homophily, providing a quick and easy way to recognise it.

Our results also show that co-author *W*-index is higher for established men and women compared to early career. There are a number of hypotheses that could explain this phenomenon. Academic communities in science remain majority men at faculty level, with the size of the majority being larger, and more static, at senior levels^22^. In most disciplines, early career scientists (postgraduate students and postdoctoral researchers) are therefore more likely to be supervised by, and consequently publishing with, men, lowering the *W*-index of all early career researchers compared to their later stage peers. Additionally, late career academics, particularly men, are more likely to hold grants^23^ that may come with funding for students and postdoctoral researchers. As these early career researchers are more likely to be women than would be expected in late stage colleagues^6^ this would increase the *W*-index of late career academics compared to early career, particularly for men as seen in the UC dataset. However, there is also evidence that high-achieving male faculty employ and train fewer women relative to other investigators^24^ which refutes this hypothesis.

The INFORMS dataset also shows an increase in *W*-index with career stage but in this case the increase is much higher for women in comparison to men. The above explanation does not explain this and partially runs contrary to it. A simpler explanation, albeit one that would be hard to confirm, is that as an individual’s career progresses, they are more able to make choices about who to work with. Women, particularly those working in very male dominated fields like those in the INFORMS dataset, may simply be choosing to work with more women. If your career feels more like a hostile obstacle course than a smooth pathway it is natural to seek out others sharing your experience.

Despite this increase with career stage, the majority of co-authors, especially for men, are still male across all but the most female dominated disciplines (Figure 1). This is a problem because scientific ideas produced by gender-diverse teams are more novel^3^ and are published in higher impact journals^20^.

A benefit of the *W*-index is its simplicity: individuals can calculate their own *W*-index, without reference to a large body of network data^25^ or unfamiliar analyses and models^26^, and can easily review their collaborator and co-author gender ratios. If we want our workforce to achieve more, people need to make choices about their own collaborations in order to support career development of all genders through all career stages. This means engaging with more diverse collaborators so that *your* lifetime *W*-index, and those close to you, all tend towards 0.5. Put bluntly, this will not happen unless everyone with influence, at any level, actively takes action and shifts teams towards gender balance at senior as well as junior levels. If equitable collaborations are not forged at scale, early career researchers will continue to fall in the hostile obstacle course of professional progression, and gender differences in other metrics of success will continue to be entrenched.

### How we recommend the *W*-index be used

#### As an author: Look at your co-author network

What proportion of your co-authors on each paper are women? Does it simply reflect the proportion of women in your field, or does it indicate a choice for more equitable working across genders? What is the largest number of men, and women, with whom you have published? *How does your collaboration network change over time, and what is the long-term average? Are you proactively inviting co-authorship with enough women researchers to shift the balance?*

#### As a collaborator: Look at your collaboration network

What is the gender balance of your longer-term colleagues compared to your trainees? What are the *W*-indices of your collaborators? *Are you fostering collaborations with women and could you do more to expand this and build more gender-diverse teams?*

#### As a group leader or member: Look at your group

How does the group’s gender make-up vary with career stage or job title? As the group leader, does your collaboration network match the diversity of the group? As a member, does the leader of your group actively provide a balanced set of role models? *Are you role-modelling or experiencing an inclusive pipeline where all genders can (and do) succeed and progress equally?*

#### As a researcher: Look at your papers’ reference lists

Do you cite women as often as men? *Do you actively seek sources from women researchers?*

#### As a speaker, or seminar organiser: Look at your colleagues

Are you organising and speaking at conferences that showcase men and women equally? Are you noticing and guiding the audience to allow diverse members to speak? *Do you actively showcase women’s work and questions?*

#### As an educator: Look at your classes and those that come before and after them

Is the *W*-index of your second-year course the same as the first-year equivalent, on average over recent years? Are women as likely to continue to further studies in your area as men? Do both genders achieve equally? *Do you actively look for ways to support and role model gender diversity for students in your classes*?

#### As a reviewer: Look at the funding proposals put forward, and those that you review

Are men in the majority of the collaborative research team, and (if so), what is the justification for this? Does the proposal include appropriate funding and planning to accommodate a gender-diverse team, and dismantle the barriers (such as access to flexible working and caring responsibilities) that disproportionately affect women? *Does your review articulate and highlight the gender-based challenges and suggest mitigation strategies?*

#### As an institutional manager: Look at your staffing and workload allocation

What gender is in the majority in senior roles, academic or managerial? Are women doing more of the caring roles (education, citizenship) and more of the internal facing research roles (departmental committees), while men take the visible, external engagement roles (grant panels, advisory and faculty boards)? *How do your retention and progression policies acknowledge, value and reward diverse contributions with respect to gender?*

#### As an institutional leader: Look at your recruitment policies

Do your hiring processes attract applications from, and then appoint, women and underrepresented minorities at least in proportion with their representation in the field? *Are you hiring women at a rate that will bring you to gender parity on a measurable time scale shorter than decades?*

#### As a person: Every day in every way

Every time you walk into a room, notice who is in the room. Notice who is talking. Notice who interrupts. *Notice who is heard*.

#### Knowledge is power, but visible action is powerful. Those who experience the problems should drive the agenda but no marginalised group should be tasked with ending their own oppression

Improving gender representation, and Equity, Diversity and Inclusion (EDI) more broadly, must be embedded into all systems and processes: “mainstreamed”, not tacked on as an additional job likely undertaken by people from under-represented minorities. This might come with short term financial risk which should be underwritten by institutions; in the longer term, diverse teams will pay dividends for impact and income^3^. An apparently meritocratic “but we hire the best people” response ignores centuries of oppression and systemic bias. The *W*-index is a simple metric that uncovers intent and action on gender equality in academic collaboration, to prompt transformative change in those systems. Think about it as the cleaning product that polishes the glass ceiling until it is crystal clear, before you smash it into tiny pieces.

## Acknowledgements

The colour scheme used in the graphics is a colour-blind friendly blend of contemporary Western gender norms (darker blue, paler pink). The authors considered using suffragette colours of purple and green but made a decision explicitly to reduce cognitive load on many readers who are accustomed to assuming red/pink=woman and blue=man. We note that these discussions added mental load and work to the manuscript writing process, which is an apt example of how making explicit, conscious decisions to be inclusive takes time. All authors should be doing, and acknowledging, this effort. A century ago, pink was considered a masculine colour, and blue was for girls^27^. Gendered colour is a social construct: we can choose to change.

The authors would like to thank Liam Gibson and James Moir for thoughtful discussions and constructive comments on the draft manuscript.

## Author Contributions

AJ and PH jointly curated and analysed the data, wrote the R code, and carried out the investigation and analysis. All authors contributed to writing the first and final drafts of the manuscript.

## Data availability

For privacy reasons, data from the University of Canterbury is not publicly available. The INFORMS dataset is downloadable with the original publication^19^. The original Maths Geneology Project dataset, including ungendered student/supervisor names is available on request at https://mathgenealogy.org/. A formatted and gendered version of the MGP data used in this paper is available in electronic supplementary information.

## Ethics Statement

This work uses data which is either fully anonymised (UC) or is publicly available. It did not require ethical approval from a human subject or animal welfare committee.

## References

1. Grogan, K. E. How the entire scientific community can confront gender bias in the workplace. Nature Ecology & Evolution 3, 3–6 (2019).

2. Huang, J., Gates, A. J., Sinatra, R. & Barabási, A.-L. Historical comparison of gender inequality in scientific careers across countries and disciplines. Proceedings of the National Academy of Sciences 117, 4609–4616 (2020).

3. Yang, Y., Tian, T. Y., Woodruff, T. K., Jones, B. F. & Uzzi, B. Gender-diverse teams produce more novel and higher-impact scientific ideas. Proceedings of the National Academy of Sciences 119, e2200841119 (2022).

4. Maas, B. et al. Women and Global South strikingly underrepresented among top- publishing ecologists. Conservation letters 14, e12797 (2021).

5. Holman, L., Stuart-Fox, D. & Hauser, C. E. The gender gap in science: How long until women are equally represented? PLoS biology 16, e2004956 (2018).

6. James, A. & Brower, A. Levers of change: using mathematical models to compare gender equity interventions in universities. Royal Society Open Science 9, 220785 (2022).

7. Berhe, A. A. et al. Scientists from historically excluded groups face a hostile obstacle course. Nature Geoscience 15, 2–4 (2022).

8. Gaule, P. & Piacentini, M. An advisor like me? Advisor gender and post-graduate careers in science. Research Policy 47, 805–813 (2018).

9. Lerchenmueller, M. J., Sorenson, O. & Jena, A. B. Gender differences in how scientists present the importance of their research: observational study. bmj 367 (2019).

10. Järvinen, M. & Mik-Meyer, N. Giving and receiving: Gendered service work in academia. Current Sociology 73, 302–320 (2025).

11. Babcock, L., Recalde, M. P., Vesterlund, L. & Weingart, L. Gender differences in accepting and receiving requests for tasks with low promotability. Am. Econ. Rev. 107, 714–747 (2017).

12. James, A., Buelow, F., Gibson, L. & Brower, A. Female-dominated disciplines have lower evaluated research quality and funding success rates, for men and women. Elife 13, RP97613 (2024).

13. Brower, A. & James, A. Research performance and age explain less than half of the gender pay gap in New Zealand universities. PLoS One 15, e0226392 (2020).

14. Gay-Antaki, M. & Liverman, D. Climate for women in climate science: Women scientists and the Intergovernmental Panel on Climate Change. Proceedings of the National Academy of Sciences 115, 2060–2065 (2018).

15. Bates, D., Mächler, M., Bolker, B. & Walker, S. Fitting linear mixed-effects models using lme4. Journal of statistical software 67, 1–48 (2015).

16. Kuznetsova, A., Brockhoff, P. B. & Christensen, R. H. lmerTest package: tests in linear mixed effects models. Journal of statistical software 82, 1–26 (2017).

17. Team, R. D. C. R: A language and environment for statistical computing. (No Title)(2010).

18. Anderson, D. & Burnham, K. Model selection and multi-model inference. Second. NY: Springer-Verlag (2004).

19. Bravo-Hermsdorff, G. et al. Gender and collaboration patterns in a temporal scientific authorship network. Applied Network Science 4, 1–17 (2019).

20. Kwiek, M. & Roszka, W. Gender-based homophily in research: A large-scale study of man-woman collaboration. Journal of Informetrics 15, 101171 (2021).

21. Suurna, M. V. & Leibbrandt, A. Underrepresented women leaders: lasting impact of gender homophily in surgical faculty networks. The Laryngoscope 132, 20–25 (2022).

22. Schoen, C., Rost, K. & Seidl, D. The influence of gender ratios on academic careers: Combining social networks with tokenism. PloS one 13, e0207337 (2018).

23. Kolev, J., Fuentes-Medel, Y. & Murray, F. Is blinded review enough? How gendered outcomes arise even under anonymous evaluation. (National Bureau of Economic Research, 2019).

24. Sheltzer, J. M. & Smith, J. C. Elite male faculty in the life sciences employ fewer women. Proceedings of the National Academy of Sciences 111, 10107–10112 (2014).

25. Avin, C. et al. in Proceedings of the 2015 Conference on Innovations in Theoretical Computer Science. 41–50 (ACM).

26. Clifton, S. M. et al. Mathematical model of gender bias and homophily in professional hierarchies. Chaos: An Interdisciplinary Journal of Nonlinear Science 29, 023135 (2019).

27. Grisard, D. “Real Men Wear Pink”? A Gender History of Color. Bright Modernity: Color, Commerce, and Consumer Culture, 77–96 (2017).

