## Supplementary Material for "The *W*-index: a novel tool to evaluate gender equity in STEM research"

Supplementary Information

1. Methods
2. Calculating *W*-index
3. *W*-index with and without the individual in question.
4. Research collaborations in a single institution (UC)
5. Co-authorship in a journal collection (INFORMS)
6. Student-supervisor relationships for research degrees (MGP)

### 1. Methods

All models were fitted in R (version 4.3.1, R Core Team 2023) using the lme4 package (Bates et al. 2015) for generalised linear mixed effects models (lmer for UC and INFORMS). The best model was selected using AIC (Burnham and Anderson 2002) and *p*-values were calculated using the Satterthwaite approximation implemented in the lmerTest package (Kuznetsova et al. 2017) as an indication of significance for fixed effect parameters in the best models. Fitting a mixed effect model to the MGP dataset (70,000 data points, 19,000 groups) was beyond the computational abilities of R on the available hardware. Instead these models were fitted in R with only fixed effects and the results were verified using a mixed effect in Matlab 2025b (fitglme).

In the MGP dataset names were gendered using first or middle names by comparing to publicly available data from the Department of Social Security, USA. It assigns a probability being female based on the proportion of names in the database. Only individuals with at least one given name that could be assigned with at least 95% accuracy were included.

### 2. Calculating *W*-index

At its core *W*-index is simply the proportion of the group who are female. The exact method of calculation is not as important as actually doing the calculation. In our examples we calculate *W*-index as follows:

*Research collaborators in a single institution*

We calculated *W*-index for each author, $w_i$, as the average proportion of female authors across all their outputs, excluding the author themselves.

$$w_i = \frac{1}{N_{outputs}} \sum \frac{Number\ of\ female\ UC\ authors\ (excl\ the\ author\ if\ female)}{Total\ number\ of\ UC\ authors - 1}$$

This method weights each output equally regardless of the number of authors. Single author outputs are not included. Outputs with many UC authors contribute to every authors *W*-index. As the calculation excludes the author in question women do not get an increase in *W*-index simply by being female! An alternative approach is to divide the total number of unique women collaborators by the total number of unique collaborators. This penalises people who publish with a single woman repeatedly alongside a larger number of men. A final version would use the total number of collaborations, i.e. non-unique, with a woman divided by the total number of (non-unique) collaborators. This would reward people who had done a single publication with a large number of women and several publications with only men. As large differences in an individual's *W*-index only appear in extreme cases choosing these alternative definitions does not change the results[1].

*Co-authorships in a journal collection*

We use similar methodology as for the UC dataset and calculate *W*-index for each unique author $i$ - publication $j$ pair.

$$w_{ij} = \frac{Number\ of\ female\ authors\ on\ publication\ j\ (excluding\ author\ i)}{(Total\ number\ of\ authors\ on\ publication\ j) - 1}.$$

Then calculate an individuals overall *W*-index as the average value across all their publications in the dataset, i.e. $w_i = \sum_j w_{ij}$.

*Student-supervisor relationships for research degrees*

*W*-index for each supervisor-student pair was simply zero or one depending on the gender of the student. A supervisor's average lifetime *W*-index was the proportion of students they had supervised who were female.

3. *W*-index with and without the individual in question

There are two ways to calculate a *W*-index for a member of a collaboration such as $w_{ij}$ for an author ($i$)-publication ($j$) pair: including or excluding the $i^{\text{th}}$ member. Including the $i^{\text{th}}$ member means that the collaboration contributes the same value ($w_{ij} = w_j$) to every member of the group. Excluding the $i^{\text{th}}$ member means that individuals take different values depending on their gender.

For example, including the individual in question a publication with two women and three men contributes $w_{ij} = 0.4$ to all authors. Excluding the $i^{\text{th}}$ member for the calculations relating this publication would contribute $w_{ij} = 0.25$ for the women authors, as each of them is working

[1] None of these methods can mitigate falsified gender diversity; for example, the 2015 story in which an astrophysicist in the USA faked a female co-author called Ursula Gamma on a number of publications and funding applications in order to improve gender diversity metrics of his group.

with one other woman, and three men. For the men authors, $w_{ij} = 0.5$ because they are working with two men and two women.

For an analytical perspective, consider a group of $N$ individuals chosen at random from a gender-diverse population with men and women equally represented. Each individual has an equal probability $p$ of being a woman or a man, independent of the gender of the other group members. Acknowledging a small proportion of nonbinary genders in the population, for this illustration let that probability be $p = 0.5$. If we calculate the *W*-index of any group member excluding the other group members it has an expected value of 0.5, i.e. the other people each have a 50% chance of being a woman.

If we calculate the *W*-index of a group member *including* themselves in the calculation for a woman the expected value is

$$W_{incl} = \frac{0.5(N-1)+1}{N},$$

i.e. the probability of each of the other $N-1$ group members being women is 0.5 but the individual in question is, by definition, a woman.

Alternatively for a man in the group we would get

$$W_{incl} = \frac{0.5(N-1)}{N},$$

as he is, by definition, not a woman. As $N \to \infty$ both these values tend to 0.5 but for smaller group sizes the woman's *W*-index is above 0.5 and the man's is below. For example, for a group of size 5, a typical number for co-authorship, her *W*-index is expected to be 0.6, whereas his is expected to be 0.4. Similarly, for a token author of one gender in a group of five, the token woman would have a *W*-index of zero excluding herself, but 0.2 including herself, while a token man would have a *W*-index of 1 excluding himself, and 0.8 including himself.

Thus, calculating *W*-index without excluding the focal individual increases the expected value for women, and lowers it for men. It is still a useful quick metric for assessing a single output; for example, a discussion panel, or a single publication. However, it is harder to interpret and support actionable behavioural change over an individual's career because the null hypothesis is not an equal *W*-index between men and women across a portfolio of outputs (e.g. lifetime publications).

4. Research collaborations in a single institution (UC)

We calculated $w_{ij}$ for each unique author ($i$)-output ($j$) pair as the number of women authors (excluding author $i$ if they were a woman), divided by the number of authors (excluding the author, regardless of gender). The *W*-index for the $i^{th}$ individual, $W_i$, is the average proportion of female authors across all their outputs, excluding the author themselves, or the mean across $j$ of all the $w_{ij}$. This method weights each output equally regardless of the number of authors, and in a population with equal numbers of men and women, who are mixing randomly and equally, we expect no gender differences between $W_i$.

We fitted linear mixed-effects models to $W_{ij}$ with fixed effects including gender, age of the author at date of publication ($Age_{ij}$), proportion of women in the field ($p_F$; estimated using department gender balance), and a random effect for author to account for multiple publications by the same author. Age at publication was calculated as $(PublicationDate - BirthYear)$ and then divided by 100 to rescale it onto a 0-1 interval, in line with the other variables in the dataset.

AIC model selection indicated that all three fixed effects feature as significant drivers of *W*-index in the best models for $W_{ij}$ (Table S1). *W*-index increases with department gender balance as expected (Table S2), but after accounting for the effects of discipline gender balance, women and older individuals have a higher *W*-index than men. The best model for $W_{ij}$ also includes a significant ($p = 0.01$) interaction for gender and age at publication such that men's W-indices increase at a greater rate with age compared to women. We also tested for the effect of a quadratic term in $p_F$ but it was rarely significant and did not score well on AIC.

**Table S1.** AIC values for models with different fixed effects, fitted to the UC dataset. All models were $w_{ij}$ ~ fixed effects + (1|*i*). The model that best describes the data (lowest AIC) is highlighted in bold.

| Fixed effects | AIC $\boldsymbol{W_{ij}}$ |
|---|---|
| $Gender * Age_{pub} * p_F$ | 10289.57 |
| $Gender + Age_{pub} * p_F$ | 10290.03 |
| $Gender * Age_{pub} + p_F$ | **10285.36** |
| $Age_{pub} + Gender * p_F$ | 10291.33 |
| $Gender + Age_{pub} + p_F$ | 10287.43 |
| $Gender * Age_{pub}$ | 10489.99 |
| $Gender * p_F$ | 10340.68 |
| $Age_{pub} * p_F$ | 10312.74 |
| $Gender + Age_{pub}$ | 10487.63 |
| $Gender + p_F$ | 10337.59 |
| $Age_{pub} + p_F$ | 10310.15 |
| $Gender$ | 10581.08 |
| $Age_{pub}$ | 10610.3 |
| $p_F$ | 10351.42 |

**Table S2.** Parameter values and significance in the best model for $\boldsymbol{W_{ij}}$ in the UC dataset.

| Fixed effect | Estimate | Std. Error | Pr(>\|t\|) |
|---|---|---|---|
| Intercept | 0.030 | 0.048 | 0.53 |
| $Gender$ (male) | -0.242 | 0.056 | 1.70E-05 |
| $Age_{pub}$ (/100) | 0.205 | 0.098 | 0.04 |
| $p_F$ (proportion of women) | 0.767 | 0.050 | 8.40E-48 |
| $Gender{:}Age_{pub}$ (male:age at publication) | 0.300 | 0.117 | 0.01063757 |

Classification: In-Confidence

For completeness, we also calculated $w_{ia}$ for each unique author ($i$)-output ($a$) pair as the number of women authors (i.e. without excluding author $i$ if they were a woman), divided by the number of all authors of known gender. The *W*-index for the $i^{th}$ individual, $W_i$, in this case is the average proportion of female authors across all their outputs, or the mean of all the $w_{ia}$. This method weights each output equally regardless of the number of authors, and all authors on a publication have the same value of $w_{ia} = w_a$ contributing to their W-index (see section 2 above). The results are qualitatively and quantitatively very similar to the results above.

5. Co-authorships in a journal collection (INFORMS)

We use the same methodology as for the UC dataset to calculate W-index $W_{ij}$ for the $i^{\text{th}}$ author using each unique author ($i$) - publication ($j$) pair.

The dataset also contains publication year, spanning more than 50 years from 1952 to 2016. Over this period the proportion of female authorships increased considerably from almost nothing to around 15%. The dataset included author gender but not age. As the data includes a unique identifier for each author we use the number of years since an author's first INFORMS publication as a proxy for career age. We used a threshold of *n* years to define early and established career authorship; i.e. authorships within *n* years of an author's first appearance in the dataset classed the author as early career for those publications, and established career for later publications. This is more likely to misclassify authors at the start of the dataset, i.e. before 1965, but, as there are far fewer publications over this period compared to later in the dataset, the effect is expected to be small.

We fitted generalised linear mixed effect models to the author-publication data $W_{ij}$, with categorical variables for gender and career stage of the author in the year of publication and a continuous variable, $p_F$, for the fraction of unique authors publishing during the year of the publication that are women. We also included a random intercept for each author to account for their publication and collaboration style. We tested thresholds of $n = 5, 10, 15$ for the career stage classification.

AIC model selection indicated that the best model for *W*-indices where authors become established after 5 or 10 years of publishing included all three fixed effects plus an interaction between career stage and gender (Table S5). Where established careers are defined as 15 years or more, the career stage effect is no longer significant, possibly because the number of author-publication pairs from established authors drops to only 13% of the data set. However, an interaction between gender and proportion of women is favoured by AIC, indicating that there are more substantial gender differences than representation alone can explain without career stage. We also tested for the effect of a quadratic term in $p_F$ but it was rarely significant and did not score well on AIC.

**Table S5** Fixed effects of models fitted to the INFORMS dataset, using 5, 10 or 15 years to define the threshold between early and established career stage. All models were $w_{ij}$ ~ fixed effects + (1|*i*). Rows highlighted in grey have no career stage fixed effect, so no difference is expected among the three columns. The best models (lowest AIC) are highlighted in bold.

| Fixed effects | AIC (5) | AIC (10) | AIC (15) |
|---|---|---|---|
| $Gender * CareerStage * p_F$ | 8343.53 | 8324.99 | 8355.71 |
| $Gender + CareerStage * p_F$ | 8357.27 | 8346.97 | 8364.26 |
| $Gender * CareerStage + p_F$ | **8336.02** | **8319.91** | 8361.03 |
| $CareerStage + Gender * p_F$ | 8345.15 | 8334.97 | 8353.98 |
| $Gender + CareerStage + p_F$ | 8351.67 | 8342 | 8361.01 |
| $Gender * CareerStage$ | 9189.49 | 9130.42 | 9217.89 |
| $Gender * p_F$ | 8343.81 | 8343.81 | **8343.81** |
| $CareerStage * p_F$ | 8636.69 | 8630.06 | 8635.51 |
| $Gender + CareerStage$ | 9201.48 | 9146.9 | 9214.72 |
| $Gender + p_F$ | 8350.7 | 8350.7 | 8350.7 |
| $CareerStage + p_F$ | 8630.92 | 8624.82 | 8632.95 |
| $Gender$ | 9282.58 | 9282.58 | 9282.58 |
| $CareerStage$ | 9649.34 | 9595.58 | 9656.33 |
| $p_F$ | 8622.51 | 8622.51 | 8622.51 |

Using the ten-year threshold, which had the lowest AIC overall, all model variables in the best model were highly significant (Table S6). The same patterns are seen in the model using a five-year threshold (Table S7). *W*-index very slowly increases as representation of women increases. After accounting for the effects of gender balance, women and later career stage individuals have a higher *W*-index, but an early career woman has a higher *W*-index than an established man, and men increase their *W*-indices more slowly with career stage than do women.

**Table S6.** Parameter values and significance in the best model for the INFORMS dataset using a ten-year threshold for reaching an established career stage.

| | Estimate | Std. Error | Pr(>\|t\|) |
|---|---|---|---|
| Intercept | 0.081 | 0.007 | 1.58E-34 |
| $Gender$ (male) | -0.079 | 0.005 | 1.08E-53 |
| $CareerStage$ (established) | 0.084 | 0.013 | 2.67E-11 |
| $p_F$ (representation of women) | 0.876 | 0.030 | 9.53E-180 |
| $Gender$:$CareerStage$ (established male) | -0.073 | 0.013 | 2.69E-08 |

**Table S7.** Parameter values and significance in the best model for the INFORMS dataset using a five-year threshold for reaching an established career stage.

| | Estimate | Std. Error | Pr(>\|t\|) |
|---|---|---|---|
| Intercept | 0.076 | 0.007 | 2.42E-30 |
| $Gender$ (male) | -0.076 | 0.005 | 5.44E-46 |

Classification: In-Confidence

| | | | |
|---|---|---|---|
| $CareerStage$ (established) | 0.057 | 0.010 | 6.53E-09 |
| $p_F$ (representation of women) | 0.888 | 0.030 | 8.23E-189 |
| $Gender$: $CareerStage$ (established male) | -0.052 | 0.010 | 5.87E-07 |

6. Student-supervisor relationships for research degrees (MGP)

We used the publicly available US Department of Social Security name-gender dataset to assign gender to students and supervisors linked to degrees from predominantly English-speaking countries (USA, UK, Australia, NZ) between 1960 and 2019. Other countries were excluded as the names could not be reliably gendered. Only names that could be assigned with over 95% accuracy were included. Despite the data being collected via citizen science methods which could easily introduce biases the proportion of postgraduate degrees awarded to women matches that given by the Association of Women in Mathematics which estimates around 21% in recent years.

After genderisation, this dataset contains unique supervisor ($i$)-student ($j$) pairs indicating the gender of supervisor and student for a relationship beginning in year $y$, such that $w_{ij} = 0$ if the student is a man, and $w_{ij} = 1$ if the student is a woman. The *W*-index for the $i^{th}$ supervisor, $W_i$, is thus the probability that that supervisor will have a woman as a student.

We assigned career stage in the same way as the INFORMS dataset: a supervisor was designated as early career for supervision relationships within ten years of their first recorded appearance as a supervisor in the dataset, and established career otherwise. The proportion of women students increases with time, so we used the proportion of women students in a given year, $p_F$, to represent their availability for supervision.

After calculating the career stage of supervisors for all supervisor-student pairs, we excluded data prior to 1960, to retain the period for which we have genderised names, and where representation of women as students and supervisors begins to increase from extremely low numbers.

We estimated the *W*-index of supervisors using a logistic regression on supervisor gender, career stage and representation of women in the student body:

$$logit(P(w_{ij} = 1)) \sim SupervisorGender * CareerStage * p_F^2.$$

As our computational abilities in R were hardware limited we did not include a random effect term for supervisor. AIC model selection indicated that the best model included all the fixed effects (Table S8), and gender, career stage, representation of women (squared), and an interaction between supervisor gender and career stage were all significant parameters (Table S9). All supervisors' *W*-indices increase with the increasing representation of women (Table S9) over the period of the dataset. Women supervisors consistently have a higher *W*-index than men but established supervisors who are men have a higher *W*-index than their early career counterparts, whereas the opposite is true for women.

We compared the findings, particularly the AIC values, with the same model including a random effect for supervisor run in Matlab 2025b which did cope with the computational task. As judged by AIC the best model with a random effect was the same as that with only fixed effects. The model estimated parameter values were very close and the final lines of best fit (shown in Figure 3) were indistinguishable in the cases with and without the random effect.

**Table S8** Fixed effects of models fitted to the MGP dataset. The best model is highlighted in bold.

| Fixed effects | AIC |
|---|---|
| $SupeGender * CareerStage * p_F^2$ | 72216.06 |
| $SupeGender + CareerStage * p_F^2$ | 72225.22 |
| $\boldsymbol{SupeGender * CareerStage + p_F^2}$ | **72208.93** |
| $CareerStage + SupeGender * p_F^2$ | 72226.15 |
| $SupeGender + CareerStage + p_F^2$ | 72223.52 |
| $SupeGender * CareerStage$ | 74547.86 |
| $SupeGender * p_F^2$ | 72268.66 |
| $CareerStage * p_F^2$ | 72545 |
| $SupeGender + CareerStage$ | 74576.26 |
| $SupeGender + p_F^2$ | 72266.08 |
| $CareerStage + p_F^2$ | 72541.98 |
| $SupeGender$ | 74753.19 |
| $CareerStage$ | 75211.87 |
| $p_F^2$ | 72561.45 |
| $SupeGender * CareerStage * p_F$ | 72275.72 |
| $SupeGender + CareerStage * p_F$ | 72290.49 |
| $\boldsymbol{SupeGender * CareerStage + p_F}$ | 72273 |
| $CareerStage + SupeGender * p_F$ | 72287.03 |
| $SupeGender + CareerStage + p_F$ | 72288.53 |
| $SupeGender * p_F$ | 72331.92 |
| $CareerStage * p_F$ | 72599.85 |
| $SupeGender + p_F$ | 72333.7 |
| $CareerStage + p_F$ | 72598.07 |
| $p_F$ | 72619.51 |

**Table S9.** Parameter values and significance in the best model for the MGP dataset.

| Parameter | Estimate | Std. Error | Pr(>\|z\|) |
|---|---|---|---|
| Intercept | -3.01437 | 0.090855 | 2.24E-241 |
| $SupeGender$ (male) | -0.65224 | 0.036015 | 2.64E-73 |
| $CareerStage$ (established) | -0.14452 | 0.070463 | 0.040262 |
| $p_F$ (representation of women) | 16.17182 | 1.091929 | 1.26E-49 |
| $p_F^2$ | -25.9732 | 3.221178 | 7.43E-16 |
| $SupeGender{:}CareerStage$ (established and male) | 0.295752 | 0.073234 | 5.38E-05 |
